# Data-informed modelling captures metabolic reprogramming and reveals branch points mediating cold stress response and growth trade-offs in rice

**DOI:** 10.64898/2026.07.07.736767

**Authors:** Fatemeh Soltani, Tiago Moreira Machado, Jan-Niklas Weder, Stefano Camborda de la Cruz, Fritz Forbang Peleke, Jedrzej Jakub Szymanski, Nadine Töpfer

## Abstract

Understanding stress-induced metabolic reprogramming in crop plants can inform breeding strategies and support the development of stress-resilient varieties. Genome-scale metabolic modelling has shown promise in elucidating network-level responses to changing environments, yet as an optimality-based approach it relies on the definition of an objective function, which is far from trivial for non-optimal conditions. To address this uncertainty, we used a time-resolved, data-informed metabolic model of rice (*Oryza sativa* L.) cold stress response as a test case, and explored two complementary approaches. We used sampling of the solution space combined with machine learning to identify reactions and pathways best characterizing the stress-induced metabolic shift, and used this information to perform Pareto analysis, placing growth and a stress-related objective in competition. This trade-off analysis identified key branch points in carbohydrate, amino acid, phenylpropanoid, nucleotide, and fatty acid biosynthesis, where resource reallocation towards stress-protection comes at the expense of growth. It further revealed differential flux modes across subcellular compartments and shifts in reducing equivalent provision as distinguishing features of the stress response. Together, these results provide a mechanistic understanding of the metabolic trade-offs and branch points governing cold stress response, and identify potential targets to optimize the cold response-growth trade-off in rice.

## Introduction

Rice (*Oryza sativa* L.) is a staple crop that constitutes the primary calorie source for more than half of the world’s population (**1**). As a crop with tropical and sub-tropical origin, rice is highly sensitive to cold stress. A drop in temperature below 15°C can adversely affect rice growth, yield, and limit its geographical spread (**2**). Early cold stress response involves initial cold perception and signal transduction (**3**) and later stages are associated with more complex metabolic adjustments (**4**) including remodelling of lipid metabolism (**5,6**) reduction in photosynthesis (**7**), accumulation of soluble sugars (**8**), and elevated levels of amino acids (**9**), particularly proline, which is a known osmoprotectant (**7**). Additionally, cold stress induces excessive production of reactive oxygen species (ROS) and of specialized metabolites such as phenylpropanoids derived compounds (**10**). These adaptive responses reflect a fundamental reallocation of limited resources from growth—to stress response-related processes, resulting in reduced growth—a phenomenon generally known as the stress-induced growth trade-off (**9**). While the processes underlying this metabolic readjustment are qualitatively well described, a quantitative analysis of the underlying metabolic trade-offs, bottlenecks and branch points requires detailed considerations of the underlying network topology and metabolic interdependencies.

Genome-scale metabolic modelling provides such a framework for quantifying how metabolic resources are allocated to meet cellular demands, by representing metabolic networks through their reaction stoichiometries. Metabolic fluxes are predicted based on known physicochemical constraints, such as nutrient availability or maximal enzyme capacities, combined with the assumption that metabolism operates towards a given objective—most commonly the maximization of biomass production. Furthermore, these models can serve as a platform for integrating omics data which yields context-specific network models capable of recapitulating metabolic states under defined conditions or across temporal scales (**12**).

Metabolic models of rice have been used to characterize metabolism across different tissues, temporal stages, and environmental conditions. Applications include elucidating seed and leaf metabolism under flooding and drought conditions (**13**), identifying essential genes during extreme photorespiration (**14**), and characterizing metabolic reprogramming in response to varying light treatments (**15**), drought (**16**), and salinity (**17**) stresses. A fundamental challenge, common to all of these examples, is the definition of an appropriate cellular objective function, as many, and potentially competing, metabolic functions are activated during the stress response. To address this, some of these studies have used alternative optima, including minimization of the sum of fluxes, minimization of the discrepancy between predicted fluxes and transcriptomics data, random sampling of the solution space, or coupling constraint-based models with crop growth models to impose physiologically informed growth constraints (**17–20**). For instance, Shaw and Cheung developed a diel metabolic model encompassing leaf, stem, and seed tissues that, when constrained by growth rates from a crop growth model, characterized day-wise, developmental stage-specific metabolic alterations under water-limited stress (**19**). Chowdhury et al. used a grain-specific model and minimized the discrepancy between predicted fluxes and transcriptomics data from control and elevated night temperature conditions to reveal key metabolic processes underlying grain chalkiness under warmer night temperatures (**18**).

Here, we challenge the assumption that a simple reduction in growth or the minimization of internal fluxes alone would adequately capture the underlying metabolic processes. Similarly, fitting fluxes to transcriptomics data is an unreliable proxy, given the generally low to moderate correlation between transcript levels and fluxes (**21**). Instead, we test unbiased and biased data-integrative modelling approaches to elucidate metabolic trade-offs, bottlenecks and branch points in metabolic stress response. To this end, we constructed a time-resolved metabolic model and contextualized it with transcriptomics data to investigate the cold-induced network-level metabolic reprogramming of rice during a 24 hour day and night cycle. To verify the general applicability of the approach, we first compared flux patterns obtained from biased and unbiased flux predictions across all time phases. In the biased scenarios we assumed the cellular objective of growth maximization and in the unbiased scenario we sampled flux patterns from the transcriptomics-data bounded solution space. In both the biased and unbiased scenario, we found that the model captures general features of early and late cold stress induced metabolic reprogramming and diurnal differences. Next, we used a machine learning pipeline on the high-dimensional flux sampling data from the unbiased scenario to identify reactions whose fluxes discriminate phases. This analysis revealed temporal patterns of metabolic reprogramming and pointed at specific reactions and pathways most strongly associated with the metabolic shift. Finally, based on these findings, we defined a more realistic cellular objective function for biased simulations and modelled the stress-induced growth trade-offs by analyzing flux patterns obtained on the Pareto frontier of two competing objectives, namely growth and osmoprotection via the accumulation of proline. This analysis revealed metabolic branch points across carbohydrate, amino acid, phenylpropanoid, nucleotide, and fatty acid metabolism that underlie the cold-response-growth trade-off in rice.

## Results

### Constructing a time-resolved model of rice cold stress response metabolism

To capture the dynamics of cold stress response in rice, we constructed a time-resolved metabolic model based on a mass- and charge-balanced genome-scale model of rice metabolism (**19**). After curating the model, we contextualized it using previously published data (**22**). This dataset comprises six time-points, control and 0.5, 2, 4, 8, 24 hours after cold initiation, and a 14 light/10 dark photoperiod (**22**). To capture the temporal dynamics of metabolic reprogramming, we replicated the model eight times, with each sub-model (or phase) representing a metabolic snapshot of a discrete time interval within the 24-hour cycle. The additional phases in the model represent two distinct dark phases. Each phase was subsequently contextualized using time-point-specific transcriptomics data using the E-Flux method (**23**). E-Flux constrains reaction fluxes based on gene expression levels, assuming that transcript abundance constrains enzyme abundance and thereby reaction flux capacity but does not fit fluxes to their corresponding transcript levels. To enable the accumulation and consumption of metabolites across phases, we connected phases via unidirectional linker reactions which enable the transfer of storage metabolites across consecutive phases as previously demonstrated (**24**). We set additional constraints to account for photorespiration by setting RuBisCO’s v_C_:v_O_-ratio to 3:1, light-dependent cellular maintenance (**25**), and day and night specific metabolic differences as in (**24**). The resulting model comprises 8 phases (phase 0 to phase 7), with 16934 reactions and 14564 metabolites, and was subsequently used to investigate the temporal cold stress-induced metabolic response in rice (Figure 1). Finally, to ensure reproducibility and broader applicability, we developed the Python package *Cobra2D*, which automates the generation of multi-tissue, time-resolved metabolic models from any user-defined constraints (see Material and Methods). All details of the model construction, the data-driven contextualization, additional constraints, and details about the simulation can be found in the Supplementary Methods.

**Figure 1:**
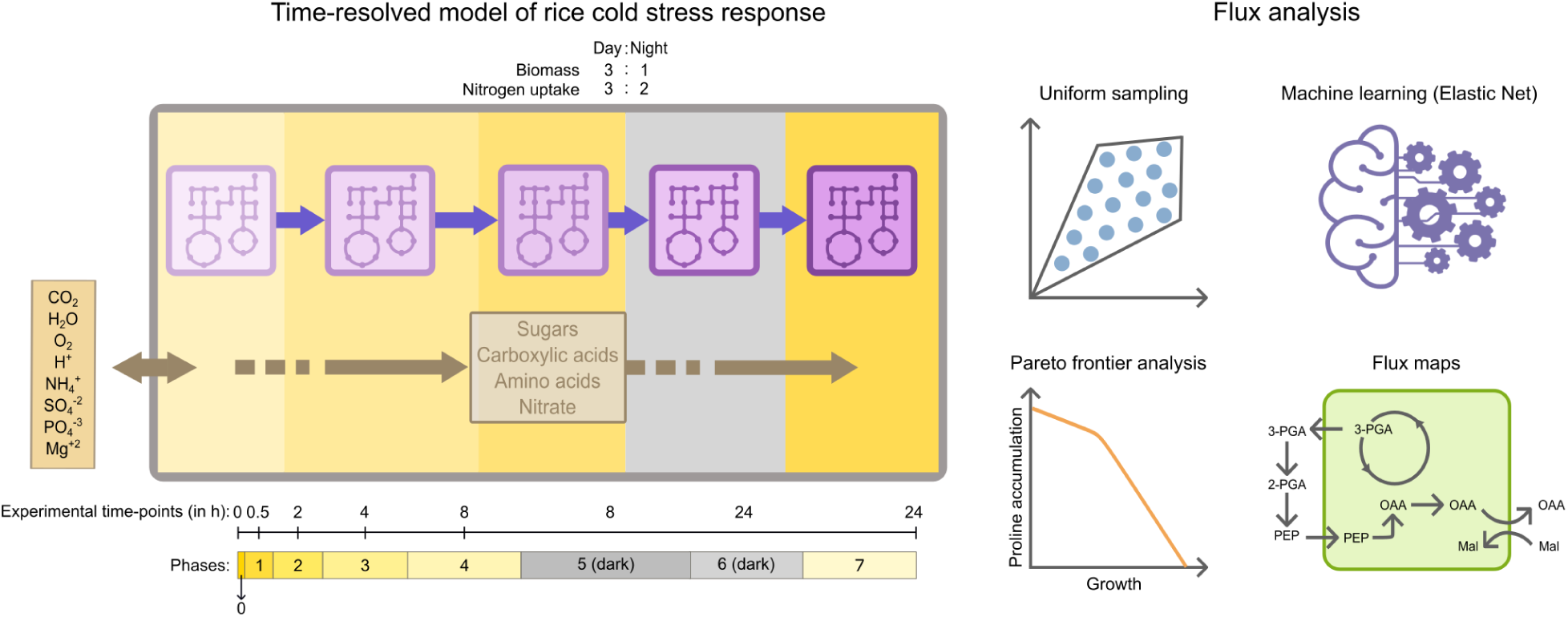
Schematic depiction of the time-resolved metabolic model of cold stress response in rice and subsequent flux analysis. Time-resolved transcriptomics data were used to generate a metabolic model of cold stress response over 24 hours. Different shades of purple depict the context-specific sub-models representing different phases informed by transcriptomics data. Purple arrows represent unidirectional linker reactions which transfer storage metabolites (beige square) between consecutive phases. The ratio of light:dark biomass production and nitrogen uptake was set as in previous studies (**24**). Experimental time-points (measured in hours after cold stress initiation) and phases in the model are depicted as a bar below. Flux patterns were obtained using biased (cellular objective-driven) and unbiased (flux sampling) approaches and subsequently analyzed.

### Analysis of flux solutions obtained from biased and unbiased flux predictions reveals distinct light–dark and early–late stress metabolic reprogramming

To obtain a general overview of the metabolic reprogramming upon cold stress induction, we compared flux patterns derived from biased and unbiased flux predictions from the transcriptomics data-bounded solution space across all phases. In the biased scenarios we assumed the cellular objective of growth maximization and used weighted parsimonious FBA (wpFBA, Supplementary Methods) to predict unique flux patterns for each of the considered phases. In contrast, in the unbiased scenario we sampled the transcriptomics data-bounded solution space without assuming any cellular objective 10000 times using optGpSampler (**26**). All data were then preprocessed before further analysis (Materials and Methods). For the biased scenario, we calculated Spearman’s Rank correlations and Euclidean distances between flux solutions. This analysis revealed strong similarity between early cold (phase 1) and control phases (phase 0), indicating minimal metabolic alterations shortly after cold initiation (Figure 2A). Intermediate phases showed progressive divergence, reflecting gradual metabolic remodelling during sustained cold exposure. The light-to-dark transition induced major metabolic shifts, with strong negative correlations and large Euclidean distances, underlining light availability as the dominant driver of divergence. Moreover, the dark phases showed strong positive correlations, indicating metabolically coherent flux patterns. The last phase exhibited a distinct profile with moderate distance to other light phases and weak to moderate correlations with other phases.

**Figure 2:**
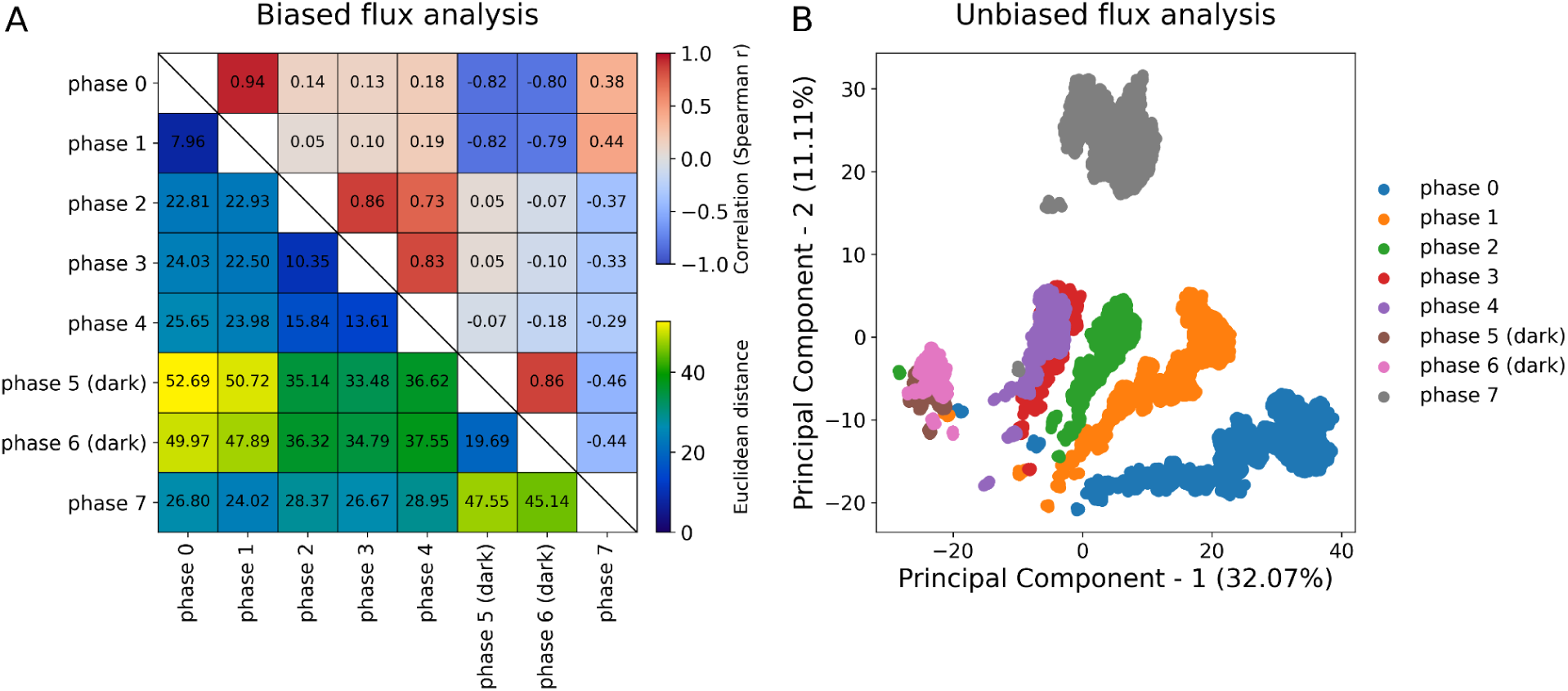
Comparative analysis of flux solutions from biased (cellular objective-driven) flux predictions and unbiased sampling of the transcriptomics data-bounded solutions space. (A) Combined matrix with Spearman’s rank correlations (upper triangle) and Euclidean distances (lower triangle) between flux solutions for each phase. (B) Principal component analysis (PCA) of metabolic flux distributions, with each point representing a flux solution colored by phases. The flux data were z-score normalized prior to the analysis in both biased and unbiased approaches (see Material and Methods).

For the unbiased approach, we performed principal component analysis (PCA) on the sampling data. This analysis revealed stress-related and day and night separation of phases with early and intermediate light phases mainly clustered together and the late cold phase (phase 7) and dark phases forming separate clusters (Figure 2B).

Together, both the biased (cellular objective-driven) and unbiased approaches demonstrate the model’s capability to capture cold stress-induced metabolic reprogramming and consistently set the late cold phase apart from other-phases—a pattern that held regardless of whether an objective function was assumed.

### Machine learning-guided flux analysis reveals temporally-distinct patterns in cold-induced metabolic reprogramming

Next, we set out to further investigate the role of individual reactions and pathways based on the unbiased flux sampling data. To identify reactions whose fluxes best discriminate between phases, we trained an Elastic Net logistic regression model (**27**) on the flux sampling data of each pair of consecutive phases and of each cold phase against control (Supplementary Methods).

Pathway enrichment analysis of reactions that best discriminate between consecutive phases revealed several well-established cold stress response pathways (Figure 3). Pathways of central metabolism including amino acid biosynthesis, photorespiration, glycolysis, TCA cycle, and photosynthesis were among the significantly enriched pathways. Oxidative stress response was consistently enriched in all consecutive phases except for the phase-5-phase-6-comparison. Purine biosynthesis and degradation pathways were among the enriched pathways, in-line with experimentally observed stress-induced purine catabolism, known to activate ABA-mediated abiotic stress responses (**28**). Phytosterol biosynthesis was enriched in the phase-2-phase-3 comparison and phase-5-phase-6 comparison, which denoted cold-induced remodelling in lipid metabolism. Proline biosynthesis, a well-characterized osmoprotectant, was enriched in both early and late phase comparisons (**29**). Results for the pathway enrichment analysis of reactions that best discriminate between individual cold phases and control yielded qualitatively similar results (Supplementary Figure 1).

**Figure 3:**
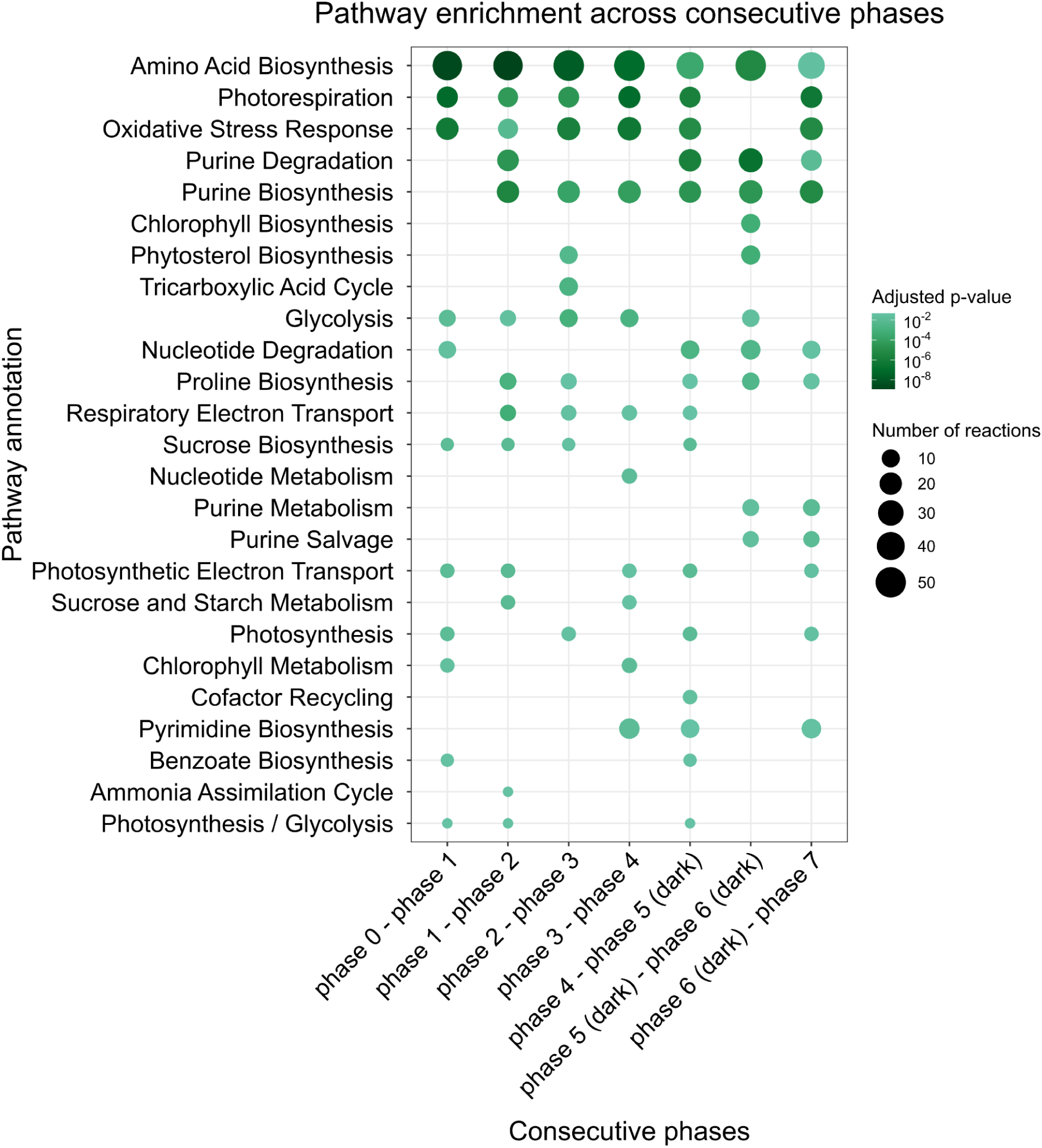
Pathway enrichment analysis across consecutive phases. The color scale represents the adjusted p-value on a -log_10_-scale and circle size denotes the number of reactions associated with a pathway. Pathways are ordered by their mean -log_10_ (adjusted p-value) across all comparisons, with pathways with the highest mean significance placed at the top of the y-axis.

The Elastic Net model identified reactions that discriminate between phases. To reveal the direction and magnitude of each reaction’s contribution to this classification, we next applied SHAP (SHapley Additive exPlanations) values (**30**) to reactions of both consecutive phases and individual cold phases versus control. This provided a biologically interpretable ranking of the discriminative reactions at each comparison (Supplementary Figure 2).

SHAP values for consecutive phases revealed that lower photosynthetic and photorespiratory fluxes were most associated with each consecutive phase comparison, except for the phase 3-phase-4 comparison in which higher photosynthetic flux discriminated the phases. Moreover, reactions involved in sucrose degradation, deoxyribonucleotide biosynthesis, CDP-diacylglycerol biosynthesis and the phenylpropanoid pathway appeared as discriminative reactions in early phases. The light-to-dark transition, was expectedly distinguished by decreased fluxes of reactions associated with photosynthesis, photorespiration, and increased fluxes of reactions associated with glycolysis, the TCA cycle, and the mitochondrial electron transport chain. Lastly, the dark-to-light phase transition was associated with increased fluxes of photosynthesis, glutamate synthesis, and the ascorbate-glutathione cycle. SHAP values of cold versus control phases revealed that reduced fluxes through reactions associated with photosynthesis, photorespiration, and glutamate biosynthesis most frequently distinguished cold from control condition (Supplementary Figure 3).

Together, these findings from the unbiased analysis of the transcriptomics data-bounded solution space pointed at specific reactions and pathways as drivers of the metabolic adjustment and reconfirmed our knowledge of cold-stress induced metabolic reprogramming.

### Accounting for proline accumulation quantifies metabolic trade-offs and branchpoints of stress response and growth

In the next step, we used the data-informed, unbiased flux sampling results to obtain a mechanistic understanding of the underlying metabolic trade-offs and branch points governing cold stress response. Among the enriched pathways identified through the unbiased approach, proline biosynthesis emerged as a suitable model pathway for examining the trade-off between growth and cold stress response as proline is a well-characterized osmoprotectant and an accumulating metabolite. In plants, proline can be synthesized via two pathways. The main route converts glutamate to proline through two enzymatic steps catalyzed by Δ^1^-pyrroline-5-carboxylate synthase (P5CS) and Δ^1^-pyrroline-5-carboxylate reductase (P5CR), consuming one ATP and two reducing equivalents, and occurs in both the plastid and the cytosol. Both plastidial and cytosolic P5CS isoforms are NADPH-dependent (**31**), while P5CR has two isoforms in both compartments which are either NADPH- or NADH-dependent (**32**). The alternative route produces proline from ornithine via ornithine aminotransferase in the mitochondria.

To study the effect of proline synthesis on metabolic fluxes, we simulated the accumulation of this osmoprotectant by introducing a proline demand reaction and performed a Pareto analysis in which we predicted metabolic flux patterns on the Pareto frontier along a gradient of decreasing growth and increasing proline accumulation (Supplementary Data 3). For our analysis we selected flux patterns at 100, 90, 50, and 10% of the maximal growth rates and refer to them as Pareto 1.0, 0.9, 0.5, and 0.1, respectively. These growth values correspond to proline content of 6.2 - 31.0 μmol m^-2^ accumulated over 24 hours and to 0, 19.2, 79.3, and 96.6% of maximal proline accumulation. These values fall well within the literature-reported range of ∼0.5 - 207.2 μmol m^-2^ for rice exposed to cold stress over the same time period (**33,34**) (Supplementary Methods and Supplementary Figure 4).

### Transcriptomics data-informed flux predictions reveal progressive reduction in photosynthesis and photorespiration during stress exposure

To investigate whether the reduction in photosynthesis, photorespiration, and glutamate synthesis identified by the SHAP value analysis, was also reflected in the Pareto-based analysis, we created fluxes maps for those pathway for both the high-biomass and high proline accumulation scenarios (Pareto 1.0 and 0.1) (Figure 4).

**Figure 4:**
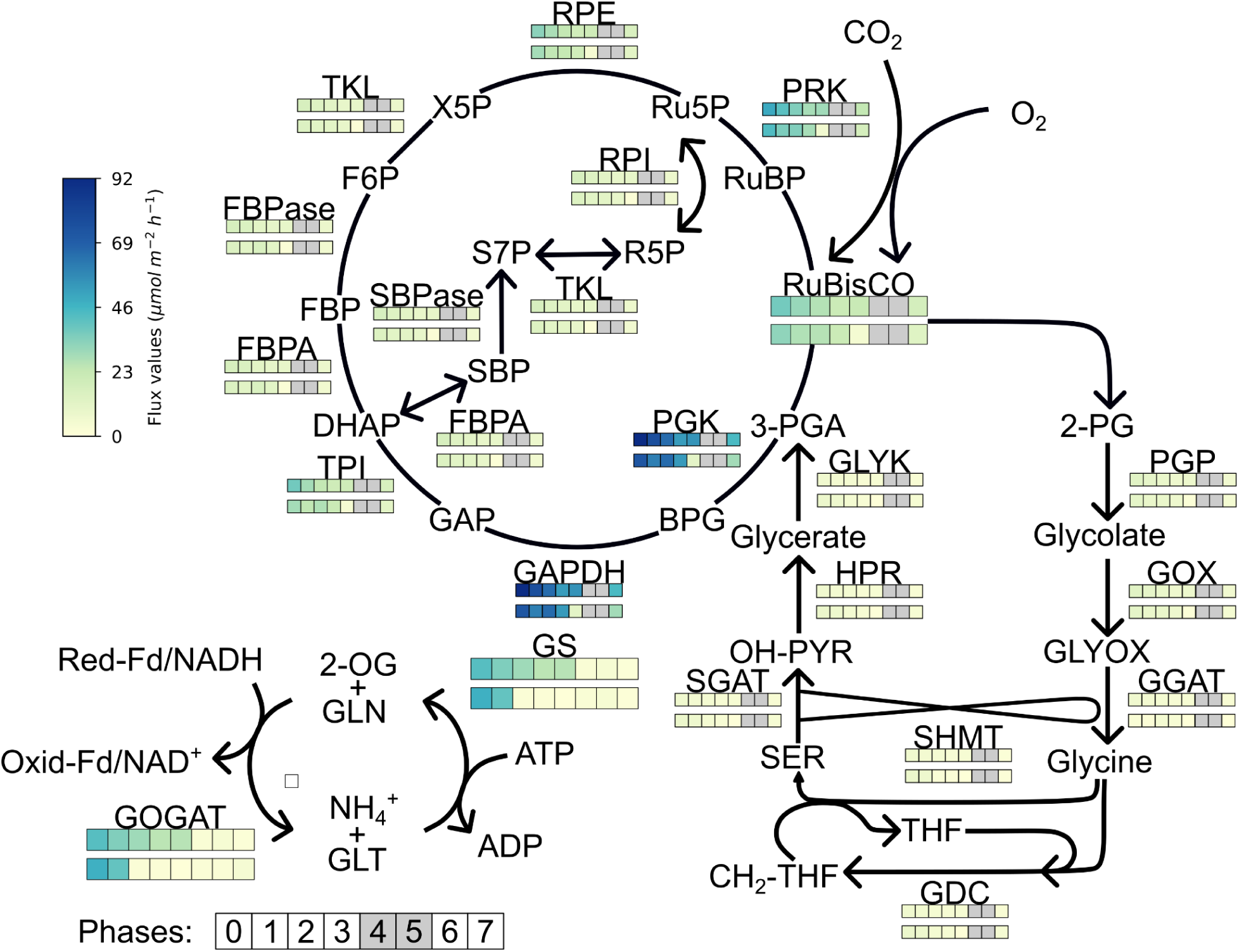
Flux-map of the Calvin-Benson-Bassham (CBB) cycle, photorespiratory, and nitrogen assimilation pathways across all phases for the high-biomass (Pareto 1.0) and high-proline accumulation (Pareto 0.1) scenarios. Fluxes of the high-biomass scenario are shown in the upper row of squares and fluxes of the high-proline accumulation scenario are depicted in the lower row, with each square denoting a single phase. For the CBB and photorespiratory cycles the dark phases (phase 5 and phase 6) are shown as grey squares. The RuBisCO v_c_:v_o_-ratio was fixed to 3:1 (**24**). Enzyme abbreviations: FBPA, fructose-1,6-bisphosphate aldolase; FBPase, fructose-1,6-bisphosphatase; GAPDH, glyceraldehyde-3-phosphate dehydrogenase; GDC, glycine decarboxylase complex; GGAT, glutamate:glyoxylate aminotransferase; GLYK, glycerate kinase; GOGAT, glutamate synthase; GOX, glycolate oxidase; GS, glutamine synthetase; HPR, hydroxypyruvate reductase; PGK, phosphoglycerate kinase; PGP, phosphoglycolate phosphatase; PRK, phosphoribulokinase; RPE, ribulose-phosphate 3-epimerase; RPI, ribose-5-phosphate isomerase; RuBisCO, ribulose-1,5-bisphosphate carboxylase; SBPase, sedoheptulose-1,7-bisphosphatase; SGAT, serine:glyoxylate aminotransferase; SHMT, serine hydroxymethyltransferase; TKL, transketolase; TPI, triosephosphate isomerase. Metabolite and cofactor abbreviations: 2-OG, 2-oxoglutarate; 2-PG, 2-phosphoglycolate; 3-PGA, 3-phosphoglycerate; BPG, 1,3-bisphosphoglycerate; CH₂-THF, 5,10-methylenetetrahydrofolate; DHAP, dihydroxyacetone phosphate; F6P, fructose-6-phosphate; FBP, fructose-1,6-bisphosphate; Fd, ferredoxin (Red-Fd, reduced; Oxid-Fd, oxidized); GAP, glyceraldehyde-3-phosphate; GLN, glutamine; GLT, glutamate; GLYOX, glyoxylate; NH₄⁺, ammonium; OH-PYR, hydroxypyruvate; R5P, ribose-5-phosphate; Ru5P, ribulose-5-phosphate; RuBP, ribulose-1,5-bisphosphate; S7P, sedoheptulose-7-phosphate; SBP, sedoheptulose-1,7-bisphosphate; SER, serine; THF, tetrahydrofolate; X5P, xylulose-5-phosphate.

In the high-biomass scenario, fluxes across all three pathways progressively decreased with prolonged cold stress duration. The high-proline accumulation scenario showed similar flux patterns in the Calvin-Benson-Bassham (CBB) and photorespiratory cycles. Only, phase 4 exhibited very low fluxes across all three pathways compared to the other light phases. Flux variability analysis revealed that this low flux reflected an alternative optima rather than a metabolic limitation (Supplementary Data 4). In contrast, nitrogen assimilation did not follow this pattern as the flux through glutamate synthase increased in the last phase to meet the demand for proline accumulation. This behaviour was specific for Pareto step 0.1, as the low- and moderate-proline accumulation scenarios showed flux patterns similar to Pareto step 1.0 (Supplementary Figure 5).

Overall, these pathway-level flux predictions revealed a coordinated metabolic response to prolonged cold stress, characterized by systematic downregulation of major anabolic processes which largely influence resource partitioning and metabolic flux modes of other central metabolic pathways across all investigated Patero steps.

### Proline accumulation reroutes CBB cycle carbon to glutamate synthesis via 3-PGA, pyruvate, and PEP branch points

Next, to investigate how increased proline production reshapes metabolism, we compared fluxes of central metabolic pathways in the last phase (phase 7) for the two contrasting scenarios of high-biomass and high-proline accumulation (Pareto steps 1.0 and 0.1). This revealed that proline accumulation relies on several metabolic branch points which mediate the stress-response and growth trade-offs. Figure 5 represents a pathway map of central carbon metabolism in which blue arrows indicate reactions with higher flux in the high-biomass scenario and red arrows indicate reactions with higher flux in the high-proline accumulation scenario.

**Figure 5.**
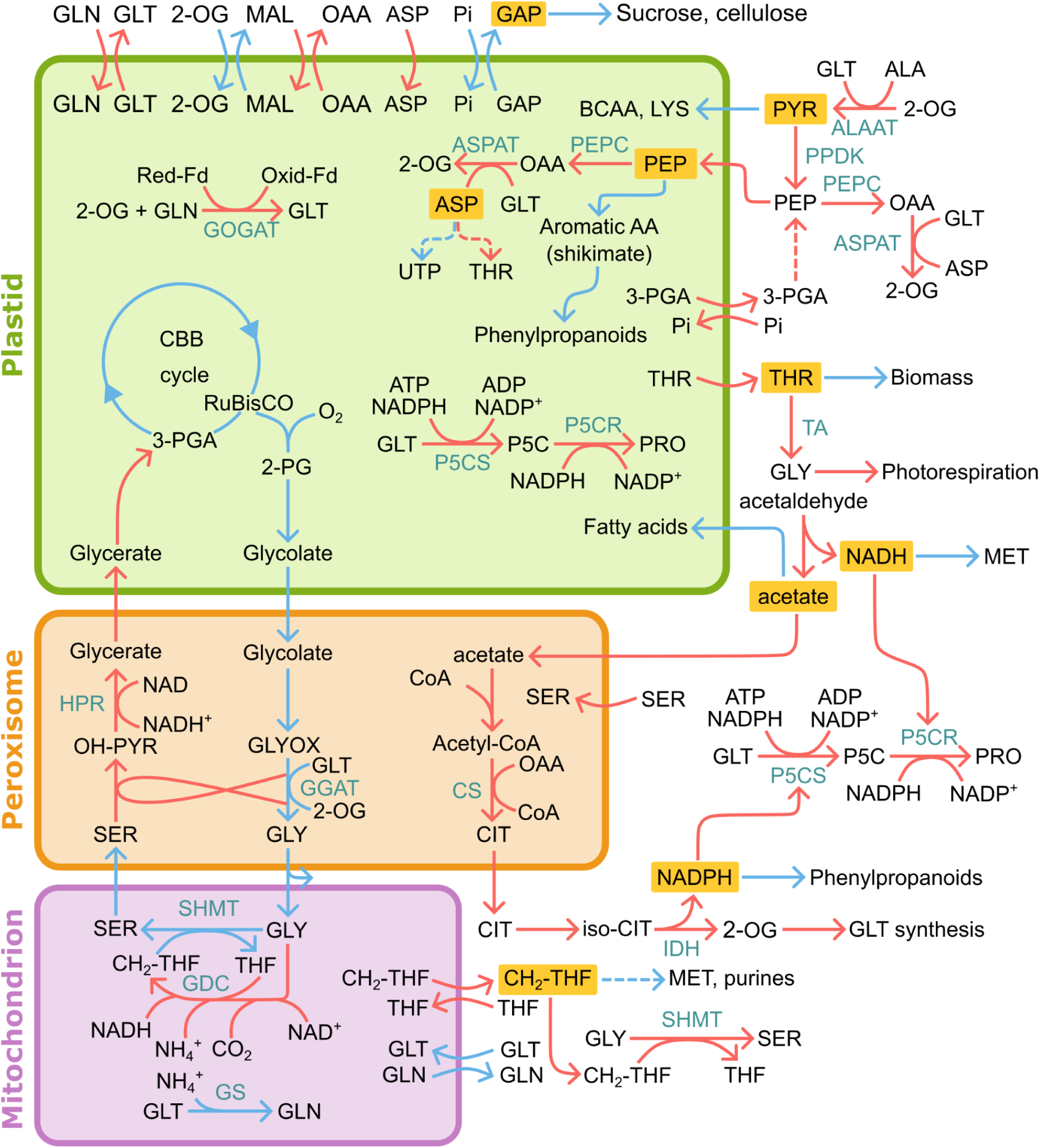
Flux map showing metabolic reprogramming of central carbon metabolism during proline accumulation. Blue arrows indicate reactions with higher flux in the high-biomass scenario (Pareto 1.0) and red arrows indicate reactions with higher flux in the high-proline accumulation scenario (Pareto 0.1) during the last phase of stress-response (phase 7). Yellow rectangles mark metabolic branch points. Enzyme abbreviations: ALAAT, alanine aminotransferase; ASPAT, aspartate aminotransferase; CS, citrate synthase; GDC, glycine decarboxylase; GGAT, glutamate:glyoxylate aminotransferase; GOGAT, glutamate synthase; GS, glutamine synthetase; HPR, hydroxypyruvate reductase; IDH, isocitrate dehydrogenase; P5CR, Δ^1^-pyrroline-5-carboxylate reductase; P5CS, Δ^1^-pyrroline-5-carboxylate synthase; PEPC, phosphoenolpyruvate carboxylase; PPDK, pyruvate orthophosphate dikinase; SHMT, serine hydroxymethyltransferase; TA, threonine aldolase. Metabolite abbreviations: 2-OG, 2-oxoglutarate; 2-PG, 2-phosphoglycolate; 3-PGA, 3-phosphoglycerate; ALA, alanine; ASP, aspartate; BCAA, branched-chain amino acids; CH₂-THF, 5,10-methylene-tetrahydrofolate; CIT, citrate; GAP, glyceraldehyde-3-phosphate; GLN, glutamine; GLT, glutamate; GLY, glycine; GLYOX, glyoxylate; ILE, isoleucine; ICT, isocitrate; LYS, lysine; MAL, malate; MET, methionine; OAA, oxaloacetate; OH-PYR, hydroxypyruvate; Oxid-Fd, oxidized ferredoxin; P5C, Δ¹-pyrroline-5-carboxylate; PEP, phosphoenolpyruvate; PHE, phenylalanine; Pi, inorganic phosphate; PRO, proline; PYR, pyruvate; Red-Fd, reduced ferredoxin; SER, serine; THF, tetrahydrofolate; THR, threonine; TRP, tryptophan; TYR, tyrosine; VAL, valine.

In the high-biomass scenario, the canonical CBB cycle operated, with glyceraldehyde-3-phosphate (GAP) directed towards ribulose-1,5-bisphosphate (RuBP) regeneration and starch synthesis in the plastid, and export to the cytosol for sucrose and cellulose synthesis, with a small fraction routed towards glycolysis. The resulting phosphoenolpyruvate (PEP) was largely imported into the plastid for aromatic amino acid synthesis and other biomass precursors via the shikimate and phenylpropanoid pathway. A smaller fraction of the cytosolic PEP was routed towards pyruvate, which was used for alanine biosynthesis via alanine aminotransferase and imported to the plastid for branched-chain amino acids and lysine biosynthesis.

The high-proline accumulation scenario was associated with lower CBB cycle activity and redirecting of 3-phosphoglycerate (3-PGA) from the CBB cycle to the cytosol. There, 3-PGA was converted to PEP via the glycolytic route, and subsequently used for glutamate and proline synthesis. This diversion increasingly limited the availability of GAP for both RuBP regeneration and the synthesis of starch and cellulose, positioning it as a branch point underlying stress-induced growth trade-offs (Supplementary Figure 6). A second metabolic branch point emerged around PEP. PEP is required both as a precursor for biomass synthesis and proline accumulation. In the high-proline accumulation scenario, PEP was mainly directed towards OAA production in the cytosol and plastid via phosphoenolpyruvate carboxylase (PEPC). OAA was then further converted to 2-oxoglutarate (2-OG) and aspartate via aspartate aminotransferase. Additionally, 2-OG was imported to the plastid through the malate/2-OG antiporter and the malate/OAA antiporter in exchange for OAA. The resulting 2-OG pool sustained glutamate, and thus proline biosynthesis. A third branch point emerged around pyruvate which was mainly used for PEP synthesis via pyruvate orthophosphate dikinase (PPDK) (which was only active in the high-proline accumulation scenario) leaving only a small fraction of pyruvate for branched-chain amino acids and lysine biosynthesis in the plastid.

These results indicate that GAP, PEP, and pyruvate act as branchpoints that support glutamate and ultimately proline biosynthesis, at the expense of biomass precursor biosynthesis.

### Photorespiratory pathway reprogramming supports carbon redirectioning from the CBB cycle

In the high-biomass scenario, we observed the canonical photorespiratory pathway fluxes. This pathway was additionally fueled by threonine produced by threonine aldolase and serine produced by serine hydroxymethyltransferase (SHMT) in the cytosol. In the high-proline accumulation scenario, consistent with reduced photosynthetic activity, photorespiratory fluxes were lower up to the reaction catalyzed by SHMT, but higher in all subsequent steps. Both the mitochondrial and cytosolic SHMT were active in each scenario, but together carried higher flux in the high-proline accumulation scenario. Interestingly, in the latter scenario, the canonical mitochondrial route in which glutamate:glyoxylate aminotransferase (GGAT) supplies glycine to mitochondrial SHMT carried lower flux while the cytosolic isoform of SHMT carried higher flux, coinciding with increased glycine production by cytosolic threonine aldolase. While the cytosolic SHMT is considered to operate in the direction of glycine to serine (**35**), thermodynamic predictions using eQuilibrator (**36**) revealed a standard Gibbs free energy (ΔG′°) of 10 kJ/mol at 4°C, thus suggesting that the reaction can proceed in both directions. This elevated downstream flux supported the increased redirection of 3-PGA towards PEP synthesis by regenerating 3-PGA in the plastid. Another metabolic branch point emerged around methylenetetrahydrofolate (CH_2_-THF), which links SHMT and glycine decarboxylase. Increased net fluxes through these reactions limited the availability of CH_2_-THF for methionine regeneration and purine nucleotides biosynthesis which serve as biomass precursors.

Together, these results position photorespiration as a key pathway supporting 3-PGA regeneration and its diversion toward PEP synthesis. We identified CH_2_-THF as a metabolic branch point mediating the stress-induced growth trade-off.

### A truncated TCA cycle provides 2-OG and reducing power required for proline synthesis

In the high-biomass scenario, asparagine accumulated during previous phases was catabolized to aspartate which fueled biomass synthesis as well as plastidial UTP production for sucrose and cellulose synthesis via UDP-glucose and threonine biosynthesis in the plastid. Threonine was exported to the cytosol and catabolized to glycine (which refueled the photorespiratory pathway) and acetaldehyde. Oxidation of acetaldehyde yielded acetate and NADH. Acetate fueled fatty acid synthesis in the plastid and 2-OG synthesis in the cytosol via peroxisomal citrate synthase and cytosolic aconitase and isocitrate dehydrogenase together forming a truncated, cross-compartmental TCA cycle.

In the high-proline accumulation scenario we observed similar flux patterns. However, here aspartate was produced by aspartate aminotransferase and not derived from stored asparagine and fluxes from aspartate to 2-OG in the truncated TCA cycle were much higher to support glutamate and ultimately proline biosynthesis. Consequently, the increased flux from aspartate to threonine reduced the flux from aspartate to UTP, thereby decreasing sucrose and cellulose biosynthesis. Similar to aspartate, redirectioning of threonine, and acetate towards acetyl-CoA for citrate synthesis came at the expense of threonine’s contribution to biomass and acetate’s contribution to fatty acid synthesis.

Notably, in both scenarios, the truncated TCA cycle used peroxisomal instead of mitochondrial citrate synthase, likely because it uses OAA generated by peroxisomal malate dehydrogenase, which produces NADH required by the photorespiratory enzyme, hydroxypyruvate reductase. In both the high-biomass and the high-proline accumulation scenario, FVA revealed that the canonical mitochondrial TCA cycle isoform can carry flux in alternative optimal solutions (Supplementary Data 4). However, fixing mitochondrial citrate synthase to its maximum FVA value rendered only the cytosolic isoforms of the downstream truncated TCA cycle enzymes—aconitase and isocitrate dehydrogenase—active, highlighting their role in supplying NADPH for proline synthesis in the cytosol. Inspection of TCA cycle fluxes during the dark phases, revealed that a canonical TCA operated in the high-biomass scenario while the truncated cycle was also active at night in the high-proline accumulation scenario (Supplementary Figures 7 and 8).

Together, this analysis indicated that during cold-stress a truncated TCA cycle reroutes aspartate-derived carbon and reducing equivalents towards proline biosynthesis. This establishes multiple branch points in fatty acid, amino acid, and UTP metabolism.

### Reducing equivalents availability determines compartmentalisation of proline biosynthesis

Given the reductive nature of the proline synthesis pathway and the photosynthetic flux remodelling observed during the stress response, we investigated how metabolism provides reducing equivalents to support increased proline biosynthesis. Plastidial NADPH was mainly used to sustain the CBB cycle across all Pareto steps. In moderate- and high-proline accumulation scenarios, an increasing proportion of cytosolic NADH and NADPH was channelled towards proline biosynthesis, diverting reducing equivalents away from the phenylpropanoid pathway and methionine regeneration (Figure 6). This suggests that dedicated cytosolic reducing power pools make the cytosol a scalable route for supporting high proline biosynthesis.

**Figure 6.**
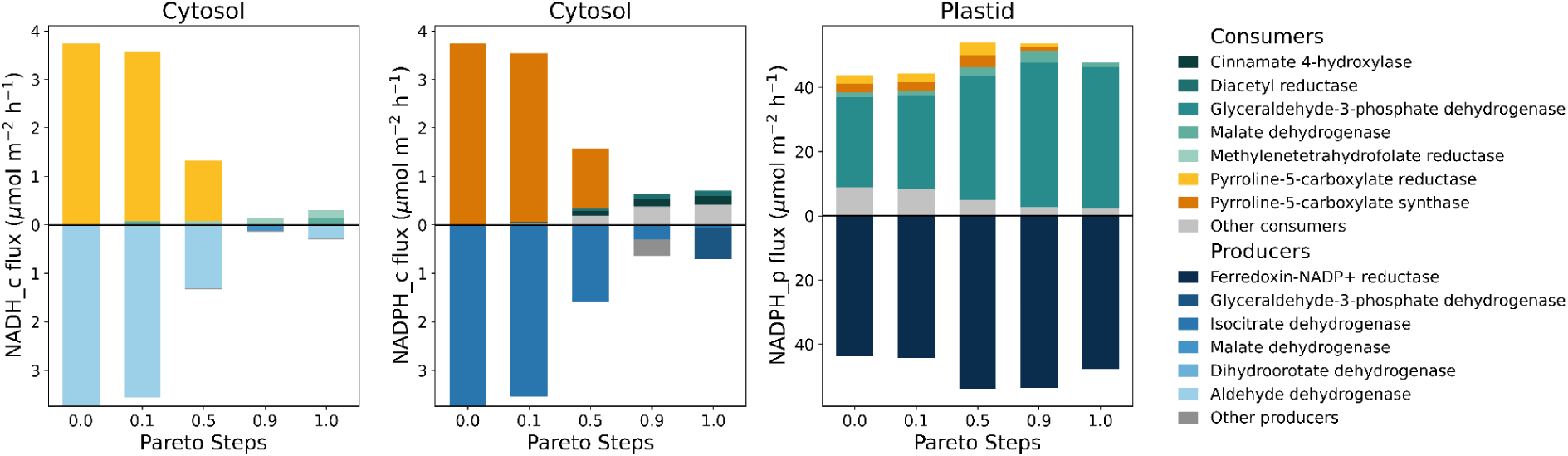
Reducing-power budget plot for the last phase across all studied Pareto steps. Shown are fluxes of NAD(P)H producing and consuming reactions in the cytosol (left and middle) and the plastid (right). Production fluxes are shown above and consumption fluxes below the zero line.

## Discussion

In this study, we investigated the choice of objective function for metabolic states under stress conditions, as cellular objectives are far from trivial to define under non-optimal conditions. To address this uncertainty, we explored two approaches. We used uniform sampling of the transcriptomics data-bounded solution space and machine learning to identify reactions, pathways, and flux deviations which best discriminate between temporal phases. This information was then used to perform Pareto analysis, which captures system behavior when growth and a stress-related objective are placed in competition. We tested this approach on a time-resolved metabolic model of rice cold stress response, using proline accumulation as a first proxy for metabolic reprogramming upon cold-stress induction. This enabled us to identify temporal cold-induced flux patterns and metabolic branch points mediating stress-induced growth-trade-offs.

### Cold-stress redirects carbon from photosynthetic end-products towards glutamate synthesis

Our model highlighted stress-induced reprogramming of starch and sucrose metabolism and predicted that reduced starch and sucrose synthesis was not only due to a reduction in photosynthesis as it has been previously reported for starch in cold-exposed wheat (**37**), and sucrose in drought-exposed rice (**38**), but also because carbon was diverted towards glutamate and proline synthesis via PEP, its conversion to OAA using PEPC, and ultimately glutamate. This diversion positions PEP as a metabolic branchpoint, because it limits its availability for the shikimate pathway and in turn phenylalanine for the phenylpropanoid pathway. The interconnection of the shikimate pathway with the CBB cycle is well-established, as at least 30% of fixed carbon feeds the shikimate pathway (**39**). However, both induction of the shikimate pathway under cold (**40**) and downregulation under drought (**41**) stresses has been reported, suggesting a stress-dependent activity of the shikimate pathway. Moreover, the association of increased PEPC activity with reduced PEP availability for aromatic amino acid synthesis has been reported for wheat transgenic lines expressing maize PEPC (**42**). In this context, we found PPDK to contribute to PEP biosynthesis in the cytosol, which is consistent with reports that PPDK accumulates in rice roots under the cold stress condition (**43**). The resulting PEP is directed towards 2-OG as a precursor for glutamate synthesis via aspartate aminotransferase. A similar route from pyruvate to glutamate and glutamine synthesis via PEP, OAA, and 2-OG via the TCA cycle has previously been reported in senescing Arabidopsis leaves, positioning PPDK as a key enzyme in this pathway (**44**). Additionally, we observed a metabolic branch point where increased redirection of pyruvate to PEP synthesis, limited pyruvate availability for branched-chain amino acids and lysine biosynthesis. Consistent with this observation, the down-regulation of amino acid biosynthetic pathways has been reported in Arabidopsis subjected to various types of abiotic stresses (**45**).

### Activity of threonine aldolase and cytosolic SHMT feeds the photorespiratory cycle

In our model, cold-stress increased fluxes through cytosolic threonine aldolase and SHMT which led to increased fluxes in the ‘returning branch’ of the photorespiratory pathway and also reduced channeling of CH_2_-THF to methionine regeneration and purine nucleotide biosynthesis. These results differ from a flux-labeling study in Arabidopsis under ambient and reduced photorespiratory conditions which reported a positive association between photorespiration and CH_2_-THF regeneration. In this study, cytosolic SHMT operated in the canonical direction and converted serine to glycine thereby generating CH_2_-THF (**35**). In Arabidopsis, the cytosolic SHMT isoform exhibits the lowest k_cat_-value for a glycine to serine conversion among all characterized isoforms, however, it has low K_m_ values for both glycine and CH_2_-THF, suggesting its high affinity for these substrates (**46**) which together with the low ΔG′° value renders the reaction to be likely reversible.

### A truncated, cross-compartmental TCA cycle provides carbon and reducing equivalents for sustained glutamate and proline synthesis

Under cold stress we observed a higher flux from aspartate to acetaldehyde, via threonine aldolase, and ultimately to the acetyl-CoA. The resulting acetyl-CoA served as a substrate for peroxisomal citrate synthase which together with cytosolic aconitase and isocitrate dehydrogenase forms a truncated, cross-compartment TCA cycle. The capacity of threonine aldolase-derived acetaldehyde to support cytosolic acetyl-CoA synthesis was demonstrated in *Saccharomyces cerevisiae*, where overexpression of threonine aldolase restored cytosolic acetyl-CoA synthesis in a pyruvate decarboxylase deficient strain (**47**). The final step of the predicted, truncated TCA cycle was catalyzed by cytosolic isocitrate dehydrogenase which generated 2-OG for glutamate synthesis and NADPH for proline synthesis. These observations are in-line with the reported contribution of cytosolic isocitrate dehydrogenase to the NADPH pool in Arabidopsis (**48**) and its role in supplying 2-OG for glutamate synthesis in pine (**49**).

### Increased demand for reducing equivalents shifts metabolic processes across cellular compartments

The Pareto analysis predicted a shift in proline synthesis from exclusive engagement of the plastid towards increasing cytosolic contribution for moderate- and high-proline accumulation scenarios. Our model suggests that this cytosolic preference is driven by the cytosol’s capacity to better meet the increased demand for reducing equivalents since plastid-derived NADPH is mainly routed towards the CBB cycle (**50**). While earlier studies suggested salinity stress-induced re-localization of P5CS1 to the plastid of Arabidopsis (**51**), later studies demonstrated cytosolic localization of both P5CS and P5CR in transgenic Arabidopsis under control and hyperosmotic conditions (**52,53**).

### Current challenges and future perspective*s*

Constraint-based metabolic modelling is a promising framework for studying stress-induced metabolic reprogramming at the network level. Besides the uncertainty in defining an appropriate objective function addressed here, other challenges should be addressed in future studies. This includes the ratio of RuBiSCO’s carboxylation:oxygenation activity, which is typically modelled as an experimentally motivated fixed ratio (**24**). Accounting for the temperature dependency of this ratio (**56**), by using a C_3_ photosynthesis biochemical model (**55**), would be a valuable refinement, once more experimental data are becoming available. Moreover, we observed that increased proline accumulation reduced starch and sucrose synthesis. Under such conditions, when the carbohydrate content is depleted, amino acids serve as alternative respiratory substrates to prevent energy depletion as reviewed previously (**56,57**). Accounting for amino acid catabolism in future modelling studies would thus enable better understanding the role of amino acid degradation for ATP generation. Furthermore, given the interconnectivity between ROS production and scavenging, the cellular redox system, and central metabolism in cold stress responses, a promising future direction would be to incorporate a ROS module with the metabolic model. This would allow stress-induced ROS accumulation to be assessed directly within the network, an approach that has already proved valuable for modelling cancer metabolism in a human model (**58**).

Finally, many amino acids do not only act as metabolites but also serve as stress signalling molecules. This balance is reflected in the antagonistic activity of TOR and SnRK1, protein kinases that sense amino acid and energy status to coordinate anabolic and catabolic responses to environmental conditions. Thus, incorporating constraint-based modelling with metabolite-orchestrated regulatory networks could enable a more direct representation of the interactions between environmental cues and metabolism.

Overall, these results provide mechanistic understanding of the metabolic branch points and compartment-specific trade-offs governing the cold stress response, and identify potential targets to optimize the cold-response–growth trade-off in rice.

## Material and Methods

### Automated reconstruction of a time-resolved large-scale metabolic model using Cobra2D

The time-resolved model reconstruction and contextualization was performed using an automated framework which we make available as a python package ‘Cobra2D’. This package streamlines the reconstruction of time-resolved and/or multi-tissue/organ models by automatically generating context-specific sub-models, adding linker reactions and/or transport reactions to connect these different sub-models, and scaling reactions to account for the lengths of the respective time intervals and subsystem sizes. Cobra2D also provides a time interval- and subsystem size-aware weighted pFBA function. The package, including detailed documentation and examples, is available via github and pip.

### Unbiased and biased flux data preprocessing

Flux data that were subjected to PCA, Elastic Net logistic regression modelling, Euclidean distance, and Spearman’s Rank correlations analyses were preprocessed by excluding non-metabolic reactions and reactions with fluxes lower than 1×10⁻⁶ across all time points. Flux values were then z-score-normalized (scikit-learn’s StandardScaler) to prevent reactions with higher absolute flux magnitudes from disproportionately influencing the results.

### Elastic Net feature selection and classification

We identified discriminating metabolic reactions between pairwise phases by training an Elastic Net logistic regression model (**27**) on the sampling data using stratified 5-fold cross-validation. We then performed hypergeometric pathway enrichment analysis (Benjamini-Hochberg FDR ≤ 0.05) on the reactions consistently selected across all cross-validation folds, and calculated SHAP values (**30**) to interpret the contribution of individual reactions to each pairwise phase classification.

### Packages

All simulations were conducted using python version 3.10.19, COBRApy python package version 0.30.0 (**59**) and the CPLEX solver (version 22.1.1.0).

## Data availability

The code needed to reproduce the results of this paper can be found at https://github.com/Toepfer-Lab/Rice_cold_response_model. The source code for Cobra2D is available at: https://github.com/Toepfer-Lab/Cobra2D.

## Author contribution

N.T. designed the research; F.S. performed the research; J.N. and S.C. developed the software package; F.S., T.M.M., F.P., J.S. and N.T. analyzed data; F.S. and N.T. wrote the article.

## Acknowledgments

Icons in figure 1 were generated using BioRender under a CC-BY License.

## Supplementary Information

**Supplementary Information:** Supplementary Methods, Supplementary Tables, and Supplementary Figures

**Supplementary Data 1:** List of manual curations applied to the rice genome-scale metabolic model.

**Supplementary Data 2:** Transcriptomics data used to contextualize the generic rice model.

**Supplementary Data 3:** Flux distributions across the Pareto frontier.

**Supplementary Data 4:** FVA results at Pareto steps 1, 0.9, 0.5, and 0.1.

**Supplementary Data 5:** Pathway enrichment analysis of reactions selected by the Elastic Net model across pairwise phase comparisons.

**Supplementary Data 6:** Pathway annotation information of model reactions.

