## Supplementary Methods for "Data-informed modelling captures metabolic reprogramming and reveals branch points mediating cold stress response and growth trade-offs in rice"

##### Reconstruction of the time-resolved metabolic model

We used an available genome-scale model of rice metabolism as our starting point **(1)**. We extensively curated the model with respect to reaction directionality, compartmentalisation, and metabolites, and added new metabolites and reactions. A full description of these modifications can be found in the Supplementary Data 1. To construct a context-specific model, we used publicly available microarray data collected from rice leaves at control (25°C) and 0.5, 2, 4, 8, and 24 hours after cold (4°C) treatment. A description of the microarray data and the experimental condition under which the data was collected can be found in Supplementary Table 1. The mean of two transcriptomics data replicates was used for model integration. We used the E-Flux algorithm **(2)** to integrate the expression data with the model using the model's Gene Protein Reaction (GPR) rules. In GPR rules which contain multiple genes, enzyme complexes should be represented by 'AND' and isozymes by 'OR' operators. Since the chosen rice genome-scale metabolic model consisted of GPRs with only 'OR' operators, we added the GPRs representing enzyme complexes from a previous rice model **(3)**.

Mapping the transcriptomics data onto these GPRs required reconciling gene identifiers between the two data sets. The model uses MSU identifiers (LOC_Os##g#####) , whereas the transcriptome data sets (Supplementary Data 2) are annotated with RAP identifiers (Os##g#######) (4,5). RAP ids were therefore converted to MSU ids using the 'riceidconvertor' R package (https://cran.r-project.org/web/packages/riceidconverter/index.html). For GPRs with 'AND' operators (enzyme complexes) the minimum transcript level and for 'OR' operators (isozymes) the sum of all transcripts were chosen to set the reaction bounds **(2)**. The resulting values were normalized based on the maximum value across all phases.

##### Defining the phases based on the experimental data setup

The duration of each phase in the model was determined such that experimental sampling time-points would fall at the mid-point of each phase (except for the dark phases which were defined based on the experiment's photoperiod). Given experimental time-points at 0 (control), 0.5, 2, 4, 8, and 24 hours after cold exposure, phase boundaries were calculated as the midpoints between consecutive time-points. For example, the boundary between the first and second phases was set at 0.25 hour (midpoint between 0 and 0.5 hour), and the boundary between the second and third phases at 1.25 hour (midpoint between 0.5 and 2 hour), yielding a second phase duration of 1 hour. The dark phase boundaries were established based on the experimental setup (the photoperiod and the time at which the dark period started and ended), with sampling initiated at 12:00 p.m. and lights switched on at 08:00 a.m. and off at 10:00 p.m. As no transcriptomic data were collected during the dark period (from 10:00 p.m. to 08:00 a.m.), we divided this 10 hour period in two phases and the border of two dark phases was placed at 16 hours (04:00 a.m.), the midpoint of 8 and 24 hours experimental time-points. The first dark phase (phase 5) was contextualized with 8 hours experimental time-points data and the second dark phase (phase 6) with 24 hours experimental time-points data. A comprehensive overview of all model phases is provided in Supplementary Table 2.

**Normalizing the flux values based on the maximum carbon dioxide uptake rate of rice**

We normalize fluxes relative to an experimentally measured $CO_2$ uptake rate of 31.5 μmol $m^2s^{-1}$ in rice **(6)**. For this purpose, we adjusted the reaction bounds by multiplying them with a scaling factor, ensuring the $CO_2$ uptake value of the control phase (phase 0) matched that of experimentally measured value.

**Accounting for the cell maintenance**

The maintenance cost of the leaf was modeled in a light-dependent manner, as previously demonstrated **(7)**. The fluxes of ATPase and NADPH oxidase reactions were set to a ratio of 3:1 according to previous measurements in Arabidopsis cell cultures **(8)**.

**Scaling the linker and biomass reaction fluxes**

To allow the transfer of storage metabolites across consecutive phases, we connected consecutive phases by adding unidirectional linker reactions. However, we did not add linker reactions that connect the last and the first phases, since these phases were in different metabolic states (control and 24 hour after cold initiation). For storage metabolites that accumulate in the vacuole, we constrained the model to account for total vacuolar storage capacity as modelled in **(7)**. A list of all storage metabolites which can be transferred between phases is given in Supplementary Table 3. The flux of all linker reactions were scaled to account for different phase lengths, as previously demonstrated **(9)**. Taking glucose as an instance, we scaled our linker reaction fluxes according to the following formulation:

$$\frac{1}{Phase\ length_{n}} Glucose\_leaf\_n \rightarrow \frac{1}{phase\ length_{n+1}} Glucose\_leaf\_n + 1$$

We modified the phase-specific biomass reactions by introducing two additional metabolites on the product side: a light- or dark-specific pseudo-biomass metabolite (light_pseudoBiomass and dark_pseudoBiomass) with a stoichiometric coefficient corresponding to the phase length which was then used for formulation of the overall_biomass reaction, and a light- or dark-specific pool biomass metabolite (light_pool_biomass and dark_pool_biomass), which was used to couple the biomass production across phases. Thus the phase-specific biomass reaction for the light phases formulated as:

$$Biomass\ precursors \rightarrow Phase\ length_{i}\ light\_pseudoBiomass + light\_pool\_biomass \quad i \in light\ phases$$

And the phase-specific biomass reaction for the dark phases formulated as:

$$Biomass\ precursors \rightarrow Phase\ length_{j}\ dark\_pseudoBiomass + dark\_pool\_biomass \quad j \in dark\ phases$$

The light- and dark-specific pool biomass metabolites were consumed by light and dark biomass constraint reactions:

$$light\_biomass\_constraint\colon\ light\_pool\_biomass \rightarrow$$

$$light\_biomass\_constraint\colon dark\_pool\_biomass \rightarrow$$

Since the light- and dark-specific pool biomass metabolites are produced by phase-specific biomass reactions and consumed only by the light or dark biomass constraint reaction, steady-state mass balance makes the flux through the biomass constraint reaction equal to the sum of the fluxes of all light- and dark-specific biomass reactions. To allow for context-specific differences in biomass production rates, we constrained our phase-specific biomass reaction fluxes to vary within a defined tolerance (±20%) around the mean flux values of all light- and dark-phase-specific biomass reactions (defined as the flux through the light and dark biomass constraint reactions divided by the number of light and dark phases). This constraint is formalized as follows:

$R_{light}$: sums the fluxes of light phase biomass reactions

$R_{dark}$: sums the fluxes of dark phase biomass reactions

$$0.8 \times \frac{R_{light}}{n_{light}} \leq v_{biomass\ i} \leq 1.2 \times \frac{R_{light}}{n_{light}} \qquad i \in light\ phases$$

$$0.8 \times \frac{R_{dark}}{n_{dark}} \leq v_{biomass\ j} \leq 1.2 \times \frac{R_{dark}}{n_{dark}} \qquad j \in dark\ phases$$

Where $R_{light}$ and $R_{dark}$ are the reactions summing up fluxes of light and dark phase biomass reactions respectively and $n_{light}$ and $n_{dark}$ are the number of light and dark phases. Finally, to enforce the 3:1 ratio of light to dark biomass production which is commonly assumed in studies modelling day and night cycles **(7)**, we introduced an overall biomass reaction that consumes the light and dark pseudo-biomass metabolites in the corresponding stoichiometric ratio:

$$overall\_biomass:\ 3\ light\_pseudo\_biomass\ +\ dark\_pseudo\_biomass \rightarrow$$

The resulting overall biomass reaction was set as the model's objective function. The code for formulation of the biomass reaction can be found in 02_analyzing_flux_solutions_without_proline.ipynb notebook.

**Setting general and day/night-specific constraints**

We modelled day and night phases by allowing the photon uptake, RuBisCO carboxylation, and cytochrome b6f activity in day-specific phases and blocking these reactions in night-specific phases. To model photorespiration, we set RuBisCO's $v_C$:$v_O$-ratio to 3:1 **(10)**. The directionality of the plastidial ATP/ADP transporter was set to only mediate the import of ATP to the plastid and its activity was restricted to the dark phase given that the knock-down mutant phenotype could only be observed in the dark condition **(11)**. We allowed the model to uptake ammonium as this is the primary nitrogen source in paddy fields **(12)**.

**Weighted parsimonious flux balance analysis (wpFBA)**

We used a modified parsimonious flux balance analysis (pFBA) that minimizes a weighted sum of fluxes, with each reaction weighted by the length of the phase.

**Elastic Net feature selection and SHAP-based interpretation**

To identify metabolic reactions that distinguish between phases, we used the data from sampling the transcriptomics data-bounded solution space to train an Elastic Net logistic regression model. Prior to model training, non-metabolic reactions, such as biomass constraint, boundary, linker reactions, and intracellular transporter reactions were excluded, such that only internal metabolic reactions were considered as features. Reaction fluxes were z-score normalized before training the model. Model performance and feature stability were evaluated using stratified 5-fold cross-validation. As Elastic Net logistic regression models select reactions that best discriminate between phases by shrinking non-informative reaction coefficients to zero, we selected reactions with non-zero coefficients in all cross-validation folds as stable discriminating features. These reactions were then subjected to pathway enrichment analysis using hypergeometric pathway enrichment analysis. This allowed us to identify significantly over-represented metabolic pathways, using the Benjamini-Hochberg false discovery rate correction (FDR ≤ 0.05) (See Supplementary Data 5 for pathway enrichment analysis of Elastic Net-selected reactions across all pairwise phase comparisons). Pathway annotations for hypergeometric pathway enrichment analysis were generated by querying the BioCyc web services **(13)** using the model's reaction identifiers. Pathway names obtained from BioCyc queries were simplified into concise, human-readable terms using a large language model (Sonnet 4.5). The original BioCyc pathway identifiers and their corresponding simplified terms are provided in Supplementary Data 6. The pathway name "cholesterol biosynthesis" was substituted with the plant-specific variant "phytosterol biosynthesis" based on BioCyc.

It is worth mentioning that we obtained classification accuracies of 1.0 for every cross-validation fold across all condition pairs, indicating that for every pairwise comparison, the Elastic Net classifier predicted the correct timepoint (label) for all test samples. While perfect accuracy on experimental data would cause scepticism, such accuracy is expected here, given the low-noise nature of model-generated flux samples. To further interpret the contribution of individual metabolic reactions to phase classifications, we calculated SHapley Additive exPlanations (SHAP) values **(14)**, which measure how much each reaction's flux contributes to the model's prediction toward one phase or the other. The SHAP values for top 20 most discriminative reactions were visualized as beeswarm plots which also show whether higher or lower flux values are associated with the selected reaction in a specific phase.

**Pareto frontier analysis and distributing proline production across phases**

To simulate cold stress-induced growth trade-offs, we modelled proline accumulation by adding a demand reaction to the last phase of the model (phase 7). To ensure the temporal distribution of proline synthesis across phases, we constrained the proline linker fluxes to lay within ±20% of the proline demand reaction flux. The linker reaction connecting the first and second phases (control and 0.5 hour after cold initiation) was excluded from this constraint, as stress-induced proline accumulation is not expected to occur under control conditions. Formally, for each constrained linker reaction flux $v_{link}^{(i)}$ the following bounds were imposed:

$$0.8 \times v_{DMpro} \le v_{link}^{(i \to i+1)} \le 1.2 \times v_{DMpro} \qquad i \in 1, \ldots, 6$$

Where $v_{link}^{(i \to i+1)}$ is the linker flux connecting phase $i$ to $i + 1$, and $v_{DMpro}$ is the flux of the proline demand reaction which was added to the last phase. We then performed Pareto analysis and placed this reaction in competition with the overall biomass reaction. The model was optimized to determine the maximum biomass flux. Then, for each Pareto step the biomass production was systematically constrained to a defined fractions of its maximum value (from a range of 100% to 0%, reduced in incremental steps of 10%), while the flux of the proline demand reaction was maximized. Flux solutions were calculated using wpFBA.

**Comparing the model-predicted proline content to experimental values**

To compare the model-predicted proline content (in µmol $m^{-2}$) over the course of 24 hours with values reported in the literature (in µmol $gFW^{-1}$ and µg $gFW^{-1}$) for the same duration, we performed a unit conversion. We used a rice specific leaf area (SLA) of 0.0417 $m^2\,g^{-1}$ as the mean of reported values in studies in different rice cultivars **(15,16)**, and a leaf dry matter content (LDMC) of 0.26 gDW $gFW^{-1}$ from Maize **(19)**, as data for rice could not be obtained. We extracted proline content values from two studies that measured proline content in rice after 24 hours of cold exposure, across different cultivars **(18)** and different seedling ages **(21)** and used the online data extraction tool WebPlotDigitizer (https://automeris.io/) to extract proline content values from the plots. The formula applied for the unit conversion, depended on the unit in which proline content was reported. For proline content values reported in µmol $gFW^{-1}$ we used the following formula to make these values consistent with the model-predicted values:

$$Proline\ content\ (\mu mol\ m^{-2})\ =\ proline\ content\ (\mu mol\ gFW^{-1})/SLA/LDMC$$

Given the extracted values for SLA and LDMC, the proline content value was 207.19 µmol $m^{-2}$ for *Oryza sativa* ssp. *indica* and 163.37 µmol $m^{-2}$ for *Oryza sativa* ssp. *Japonica* cultivars. For proline content values reported in µg $mg^{-1}$FW we used the following formula to convert the values to flux values:

$$Proline\ content\ (\mu mol\ m^{-2})\ =\ proline\ content\ (\mu g\ gFW^{-1})/MW/SLA/LDMC$$

where MW is the molar weight of proline which is 115.13 g $mol^{-1}$. Using the literature-based values for SLA and LDMC results in proline content values between 0.51 - 1.17 µmol $m^{-2}$.

### Supplementary Tables

#### Supplementary Table 1. Features of the integrated microarray data.

| Accession number | Rice cultivar | Plant age and growth stage | Temperature | Photoperiod | Time-points | Number of mapped genes | Reference |
|---|---|---|---|---|---|---|---|
| E-MEXP-37 18 | Jumli Marshi (japonica) | three weeks old (vegetative) | 25°C/20°C for control and 4°C for cold | 14 Light/10Dark | Control and 0,5, 2, 4, 8, 24 hours after cold exposure | 2034 out of 2430 model genes | (20) |

#### Supplementary Table 2. Detailed description of model phases.

| Phase ID | Length (h) | light / dark | Phase boundaries (h) | Clock time | Data integration |
|---|---|---|---|---|---|
| leaf-0 | 0.25 | Light | 0 - 0.25 | 12 - 12:15 p.m | Control |
| leaf-1 | 1 | Light | 0.25 - 1.25 | 12: 15 - 13:15 | 0.5 h after cold |
| leaf-2 | 1.75 | Light | 1.25 - 3 | 13:15 - 15:00 | 2 h after cold |
| leaf-3 | 3 | Light | 3 - 6 | 15 - 18 | 4 h after cold |
| leaf-4 | 4 | Light | 6 - 10 | 18 - 22 | 8 h after cold |
| leaf-5 | 6 | Dark | 10 - 16 | 22 - 4 | 8 h after cold |
| leaf-6 | 4 | Dark | 16 - 20 | 4 - 8 | 24 h after cold |
| leaf-7 | 4 | Light | 20 -24 | 8 - 12 p.m | 24 h after cold |

#### Supplementary Table 3. List of storage metabolites allowed to accumulate across phases.

| Type | Name | |
|---|---|---|
| Sugars | Starch | Fructose |
| | Glucose | Sucrose |
| Carboxylic acids | Malate | Citrate |
| | Fumarate | |
| Amino acids | Proline | Cysteine |
| | Tyrosine | Glutamate |
| | Asparagine | Glutamine |
| | Serine | Glycine |

| | | |
|---|---|---|
| | Methionine | Tryptophan |
| | Valine | Isoleucine |
| | Leucine | Alanine |
| | Phenylalanine | Arginine |
| | Thereonine | Aspartate |
| | Lysine | Gamma-aminobutyric acid |
| | Histidine | |
| Nitrogen | Nitrate | |

#### Supplementary Figures

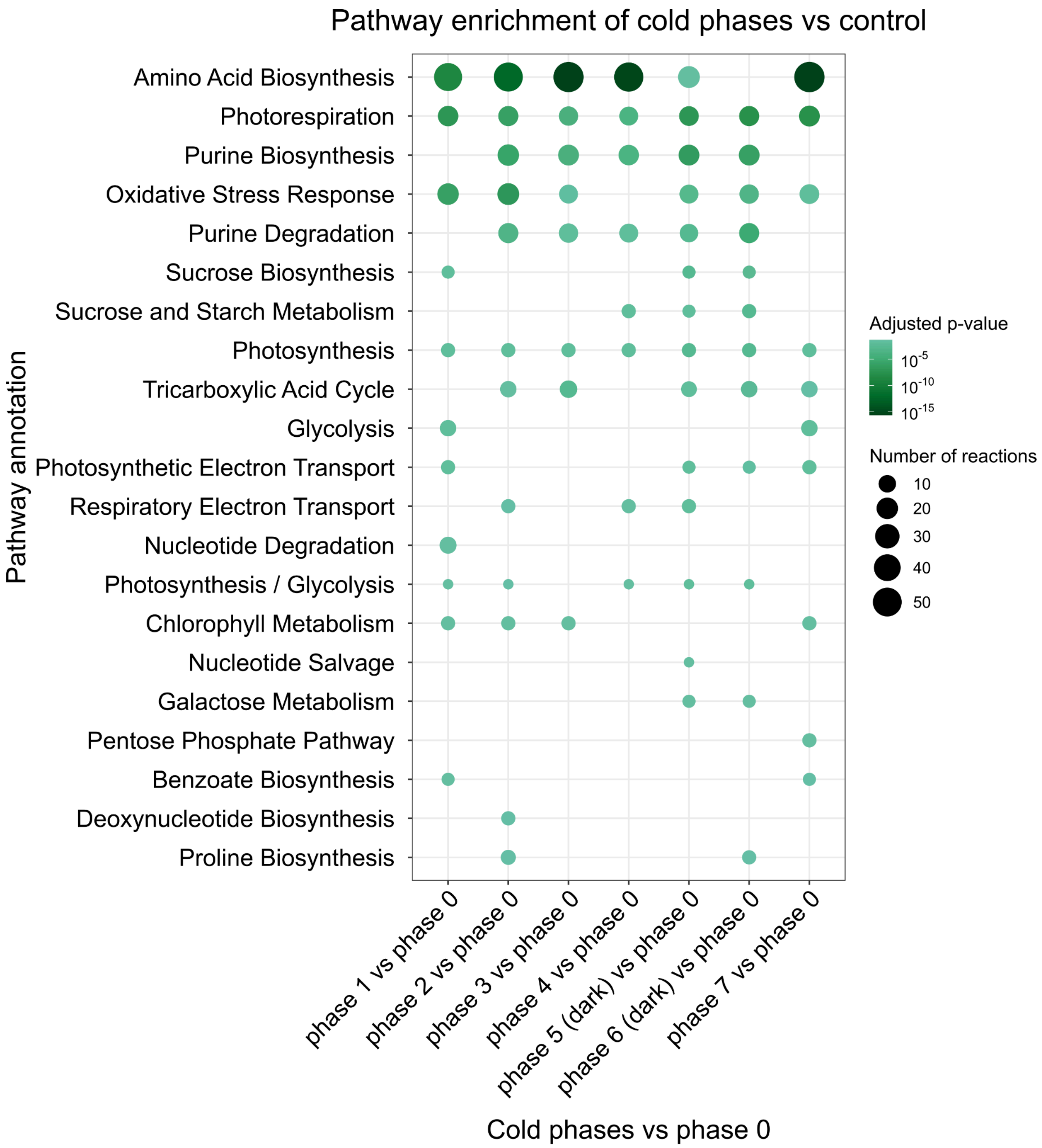


**Supplementary Figure 1. Pathway enrichment analysis for cold phases versus control.** The color scale represents the adjusted p-value on a $-\log_{10}$-scale and circle size denotes the number of reactions associated with a pathway. Pathways are ordered by their mean $-\log_{10}$ (adjusted p-value) across all comparisons, with pathways with the highest mean significance placed at the top of the y-axis.

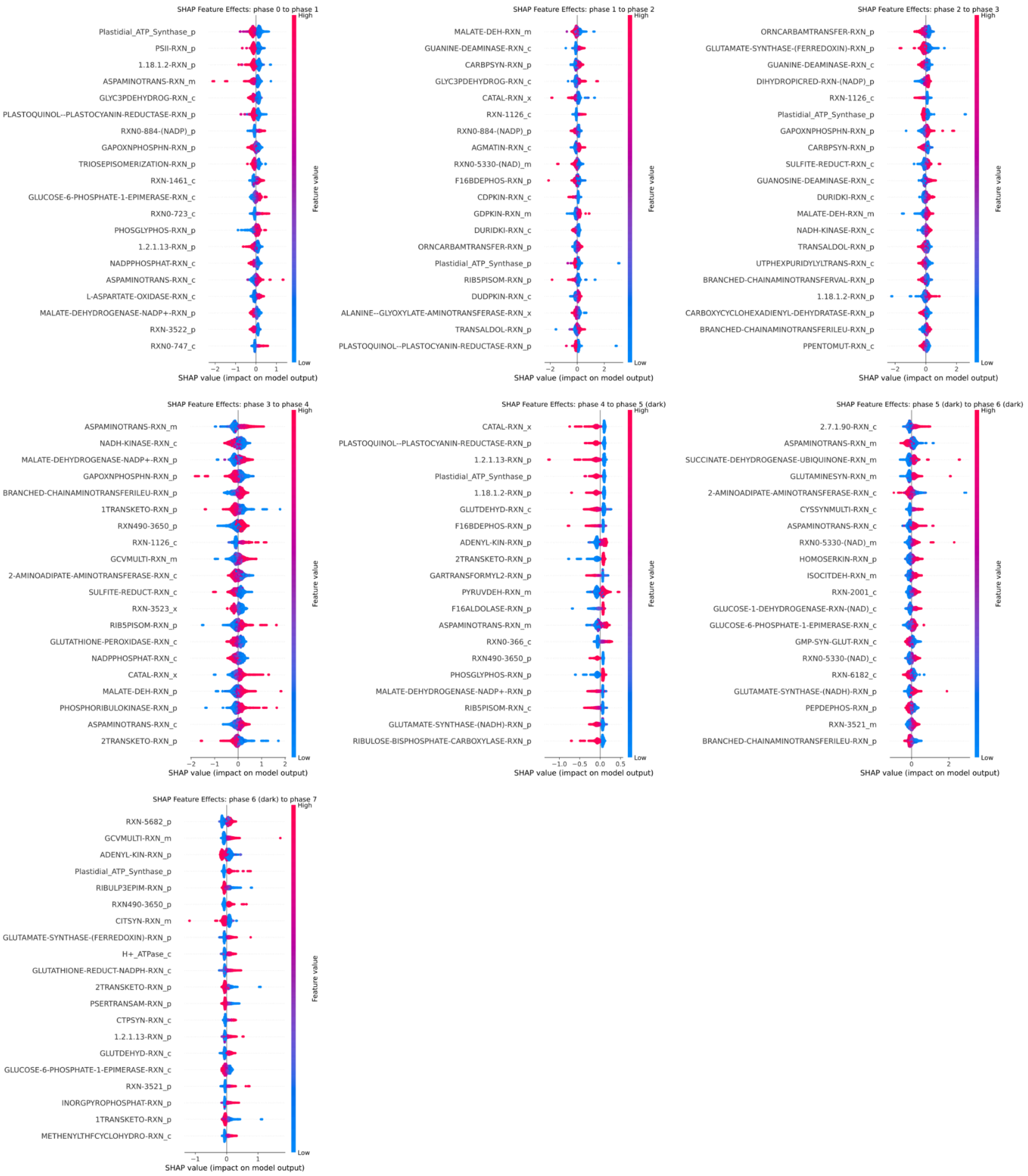


**Supplementary Figure 2. SHAP (SHapley Additive exPlanations) beeswarm plot for consecutive phases, reflecting the magnitude and direction of each reaction's contribution to the Elastic Net model's prediction.** Each dot represents a single sampled flux solution, the dot color indicates the reaction's flux value in that sampled flux solution. Top 20 reactions ranked by mean absolute SHAP value are shown.

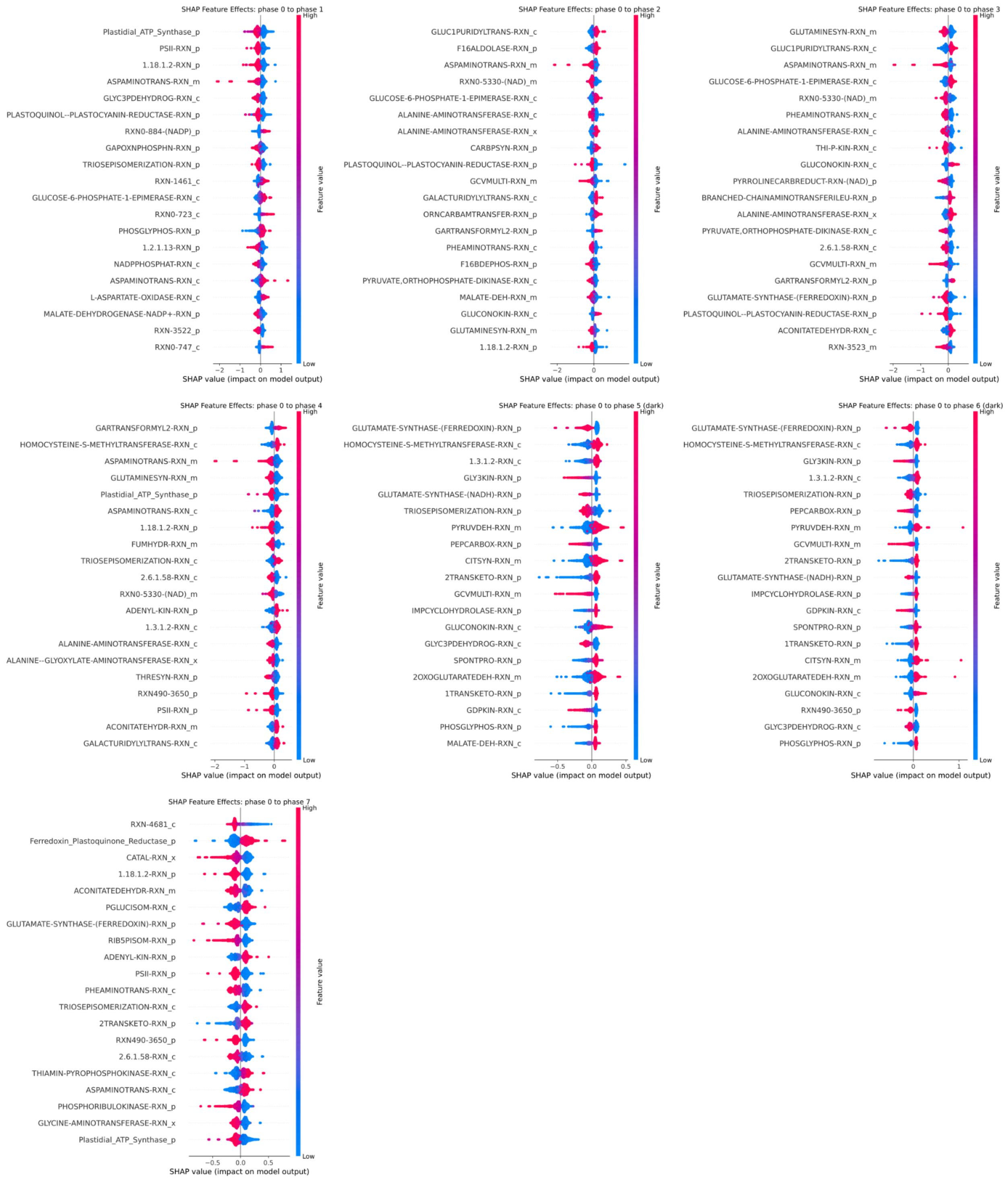


**Supplementary Figure 3. SHAP (SHapley Additive exPlanations) beeswarm plot for cold phases and control, reflecting the magnitude and direction of each reaction's contribution to the Elastic Net model's prediction.** Each dot represents a single flux sampling solution, with dot color indicating the reaction's flux value. Top 20 reactions ranked by mean absolute SHAP value are shown.

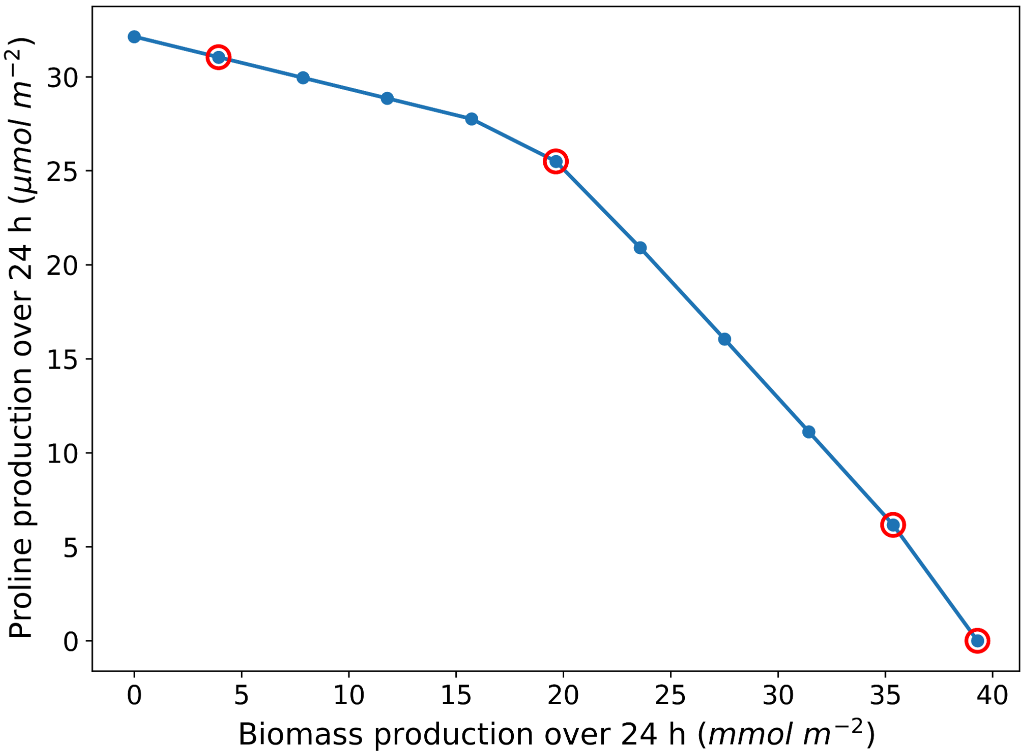


**Supplementary Figure 4. Pareto frontier representing the change in growth (measured as biomass production) versus proline content over the course of 24 hours after stress application.** The blue line shows the change in biomass production with respect to proline content with each point representing different pareto steps from step 0 (0% growth) to step 1 (100% growth) increment in 0.1 steps. The points highlighted with red circles denote selected Pareto steps (0.1, 0.5, 0.9, and 1.0) for further flux analyses.

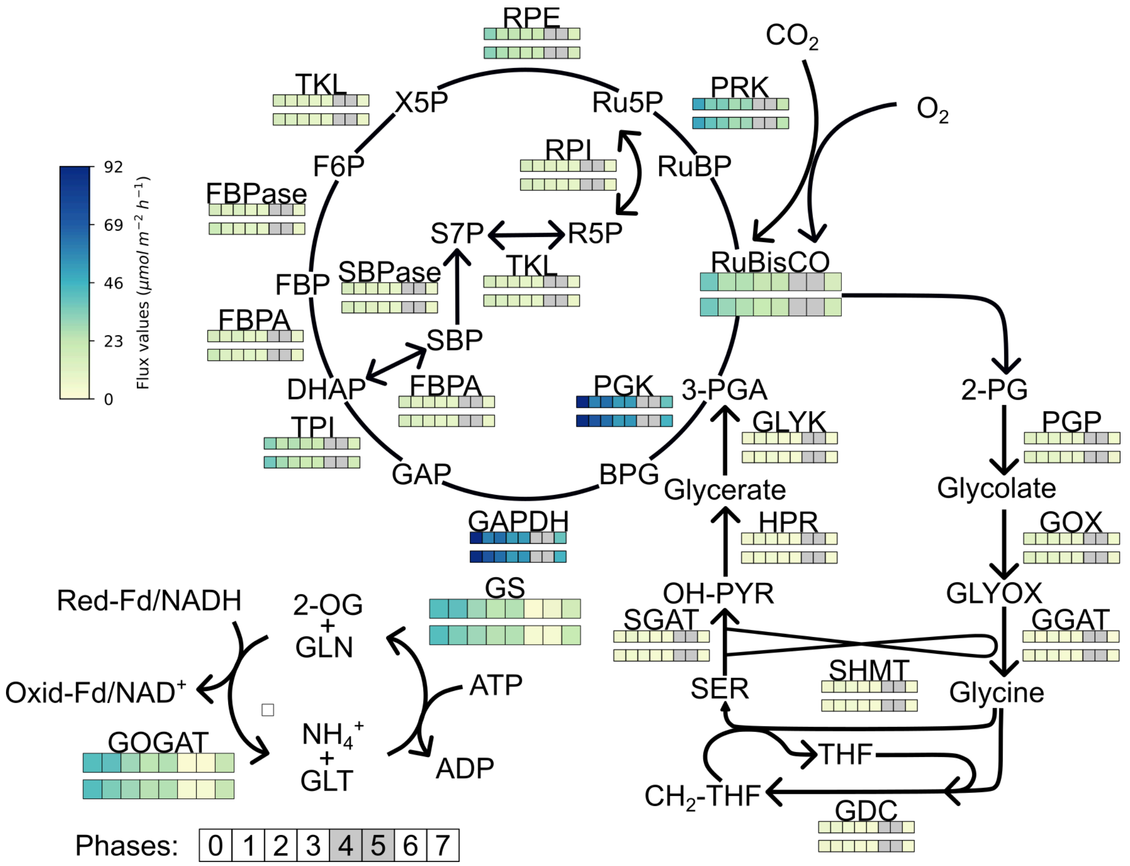


**Supplementary Figure 5. Flux-map of the Calvin-Benson-Bassham (CBB) cycle, photorespiratory, and nitrogen assimilation pathways across all phases in the low- (Pareto 0.9) and moderate- (Pareto 0.5) proline accumulation scenarios.** Fluxes of the low-proline accumulation scenario are shown in the upper row of squares and fluxes of the moderate-proline accumulation scenario are depicted in the lower row, with each square denoting a single phase. For the CBB and photorespiratory cycles the dark phases (phase 5 and phase 6) are shown as grey squares. The RuBisCO $v_c$:$v_o$-ratio was fixed to 3:1 **(10).** Enzyme abbreviations: FBPA, fructose-1,6-bisphosphate aldolase; FBPase, fructose-1,6-bisphosphatase; GAPDH, glyceraldehyde-3-phosphate dehydrogenase; GDC, glycine decarboxylase complex; GGAT, glutamate:glyoxylate aminotransferase; GLYK, glycerate kinase; GOGAT, glutamate synthase; GOX, glycolate oxidase; GS, glutamine synthetase; HPR, hydroxypyruvate reductase; PGK, phosphoglycerate kinase; PGP, phosphoglycolate phosphatase; PRK, phosphoribulokinase; RPE, ribulose-phosphate 3-epimerase; RPI, ribose-5-phosphate isomerase; RuBisCO, ribulose-1,5-bisphosphate carboxylase; SBPase, sedoheptulose-1,7-bisphosphatase; SGAT, serine:glyoxylate aminotransferase; SHMT, serine hydroxymethyltransferase; TKL, transketolase; TPI, triosephosphate isomerase. Metabolite and cofactor abbreviations: 2-OG, 2-oxoglutarate; 2-PG, 2-phosphoglycolate; 3-PGA, 3-phosphoglycerate; BPG, 1,3-bisphosphoglycerate; $CH_2$-THF, 5,10-methylenetetrahydrofolate; DHAP, dihydroxyacetone phosphate; F6P, fructose-6-phosphate; FBP, fructose-1,6-bisphosphate; Fd, ferredoxin (Red-Fd, reduced; Oxid-Fd, oxidized); GAP, glyceraldehyde-3-phosphate; GLN, glutamine; GLT, glutamate; GLYOX, glyoxylate; $NH_4^+$, ammonium; OH-PYR, hydroxypyruvate; R5P, ribose-5-phosphate; Ru5P, ribulose-5-phosphate; RuBP, ribulose-1,5-bisphosphate; S7P, sedoheptulose-7-phosphate; SBP, sedoheptulose-1,7-bisphosphate; SER, serine; THF, tetrahydrofolate; X5P, xylulose-5-phosphate**.**

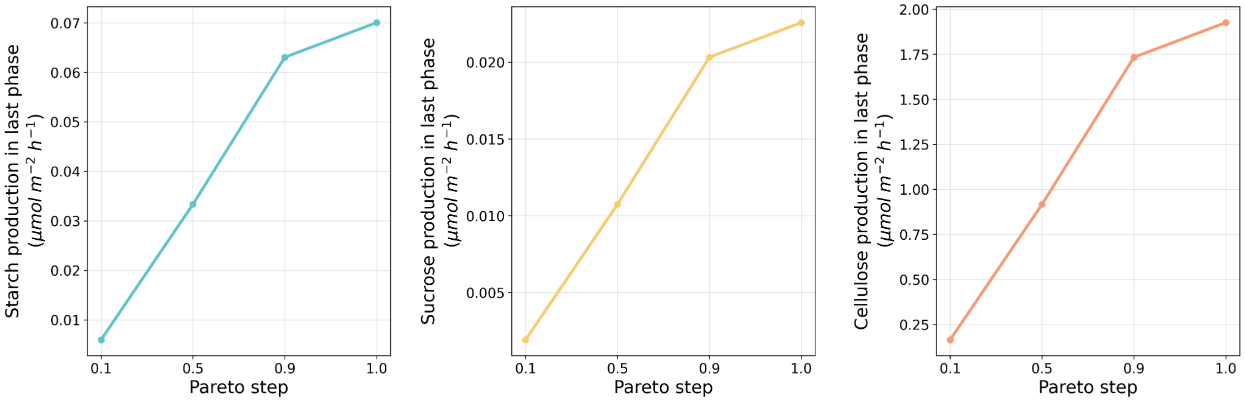


**Supplementary Figure 6. Carbohydrates synthesis fluxes in the last phase for selected Pareto steps.** From left to right flux of starch, sucrose, and cellulose synthesis for Pareto steps 0.1, 0.5, 0.9, and 1.0.

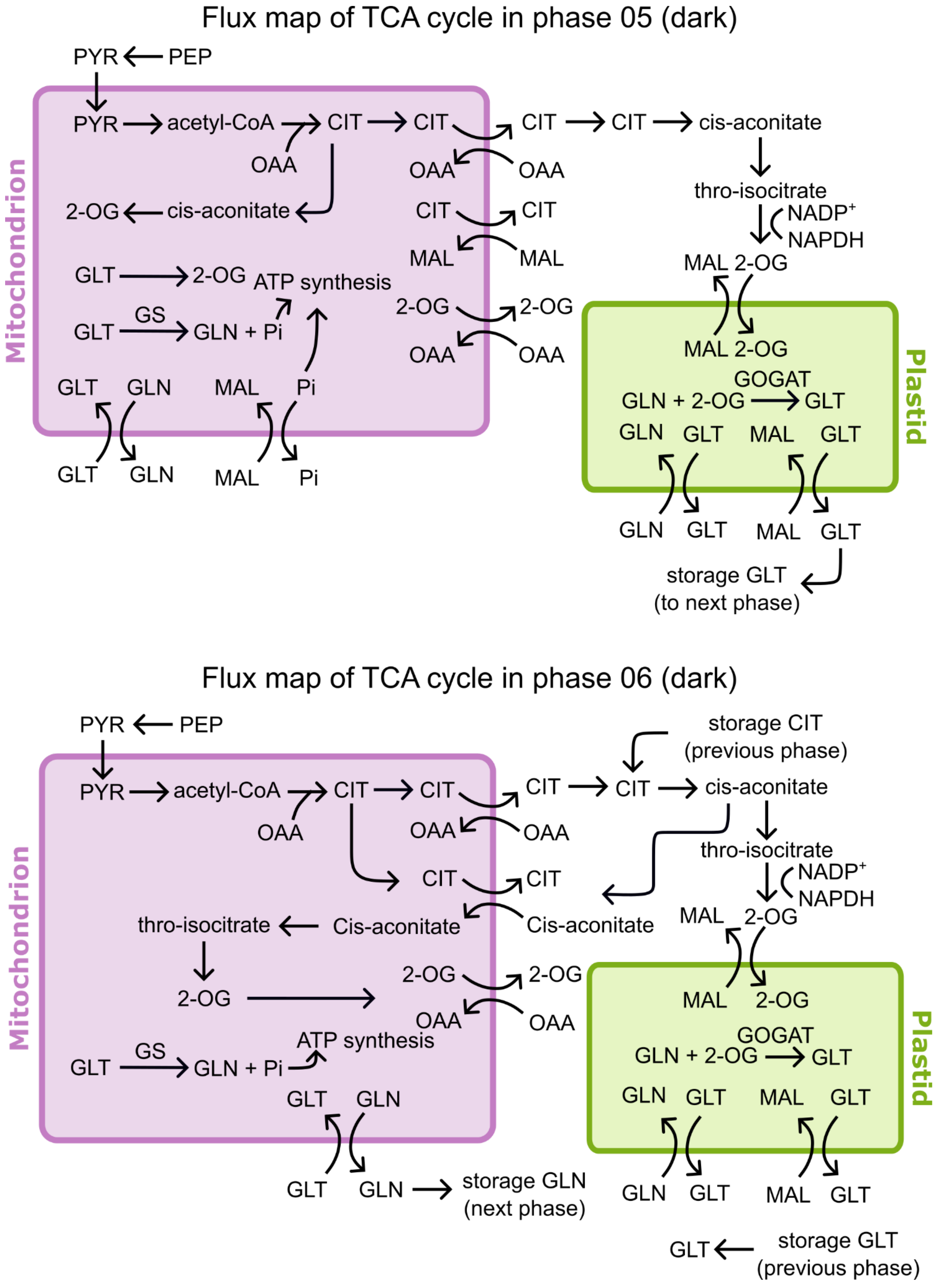


**Supplementary Figure 7. Flux modes of the truncated TCA cycle and nitrogen assimilation pathway during the dark phases (phase 5 and phase 6) in the high-proline accumulation scenario (Pareto 0.1).**

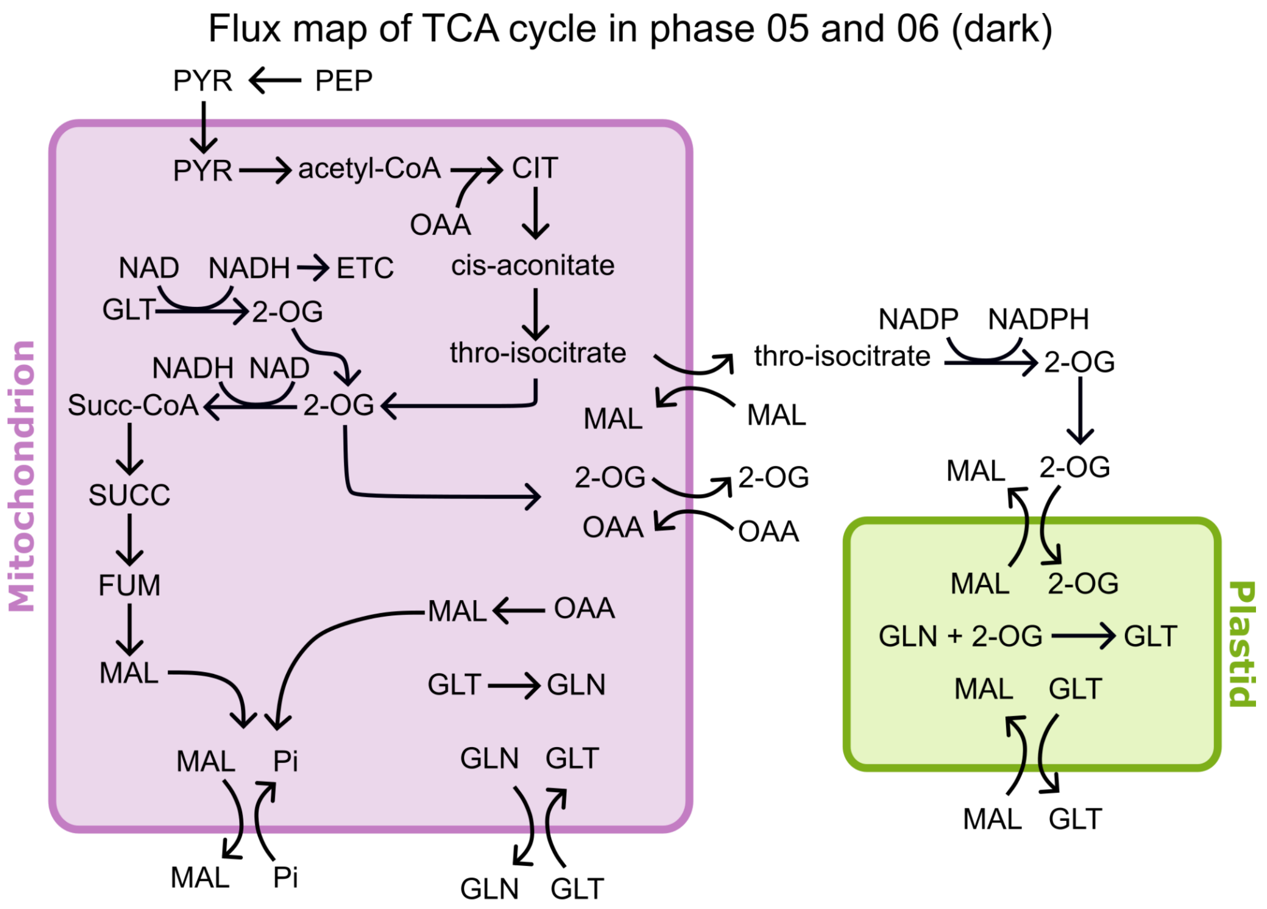


**Supplementary Figure 8. Flux modes of the truncated TCA cycle and nitrogen assimilation pathway during the dark phase (phase 5 and phase 6) in the high-biomass scenario (Pareto 1.0).**
